# Sexual difference in defense can drive the evolution of imperfect Müllerian mimicry in the less defended sex

**DOI:** 10.64898/2026.05.21.727037

**Authors:** Chi-Yun Kuo

## Abstract

Müllerian mimicry is the convergent evolution of warning signals among sympatric prey driven by predator learning. Theory therefore predicts signal homogeneity both within communities and within species that participate in Müllerian mimicry. Though rare, sexual dimorphism in Müllerian species does occur, but the underlying eco-evolutionary mechanisms are still relatively unexplored. Basing on the biology of aposematic butterflies, this study uses a modeling approach to test the hypothesis that sexual difference in defense can lead to the evolution of imperfect Müllerian mimicry in the less defended females as the consequence of opposing demands to minimize the cost of automimicry while maximizing reproductive output. Additionally, both the occurrence and degree of sexual dimorphism would decrease when the less defended sex becomes more valuable for reproduction, for example when offspring sex ratio is male biased or when females can mate only once in their lifetime. Findings from this study could help explain the evolution of extreme sexual dimorphism in some Müllerian systems, in which each sex mimics different models. Moreover, through understanding this intriguing exception to the rule, we will be able to gain a more complete picture of how a multitude of selective forces might shape the diversity in prey phenotypes.

## Introduction

Müllerian mimicry is the convergent evolution of warning signals among sympatric prey targeted by the same predators [1–3]. Such convergence in signal appearance is driven by positive frequency-dependent selection due to number-dependent avoidance learning — each predator must sample a fixed number of aposematic prey to develop avoidance [1,4] (but see [5,6]). We therefore would expect signal homogeneity both within communities and within species that participate in Müllerian mimicry. However, this expectation is not always met in natural communities. For example, it is not uncommon to observe the coexistence of multiple distinct warning signals within communities [7–9]. Furthermore, there can be sexual dimorphism in color patterns within Müllerian mimetic species, such as *Dilophotes* beetles [10,11], *Oophaga* poison frogs [12], and *Euploea* butterflies (Figure 1). Even though sexual dimorphism in mimetic systems does occur, the majority of the cases are sex-limited Batesian mimicry, in which only one sex of an undefended species, or the undefended sex within an overall defended species, mimics a sympatric model [13–16]. The selective pressures driving sex-limited Batesian mimicry in these cases include avoiding costly sexual interactions with the model species, developmental constraint coupled with sexually contrasted predation, and mitigating competition with the model species [17,18]. However, the discrepancy between expected signal monomorphism within species and observed sexual dimorphism in strictly Müllerian systems is yet to be reconciled.

**Figure 1.**
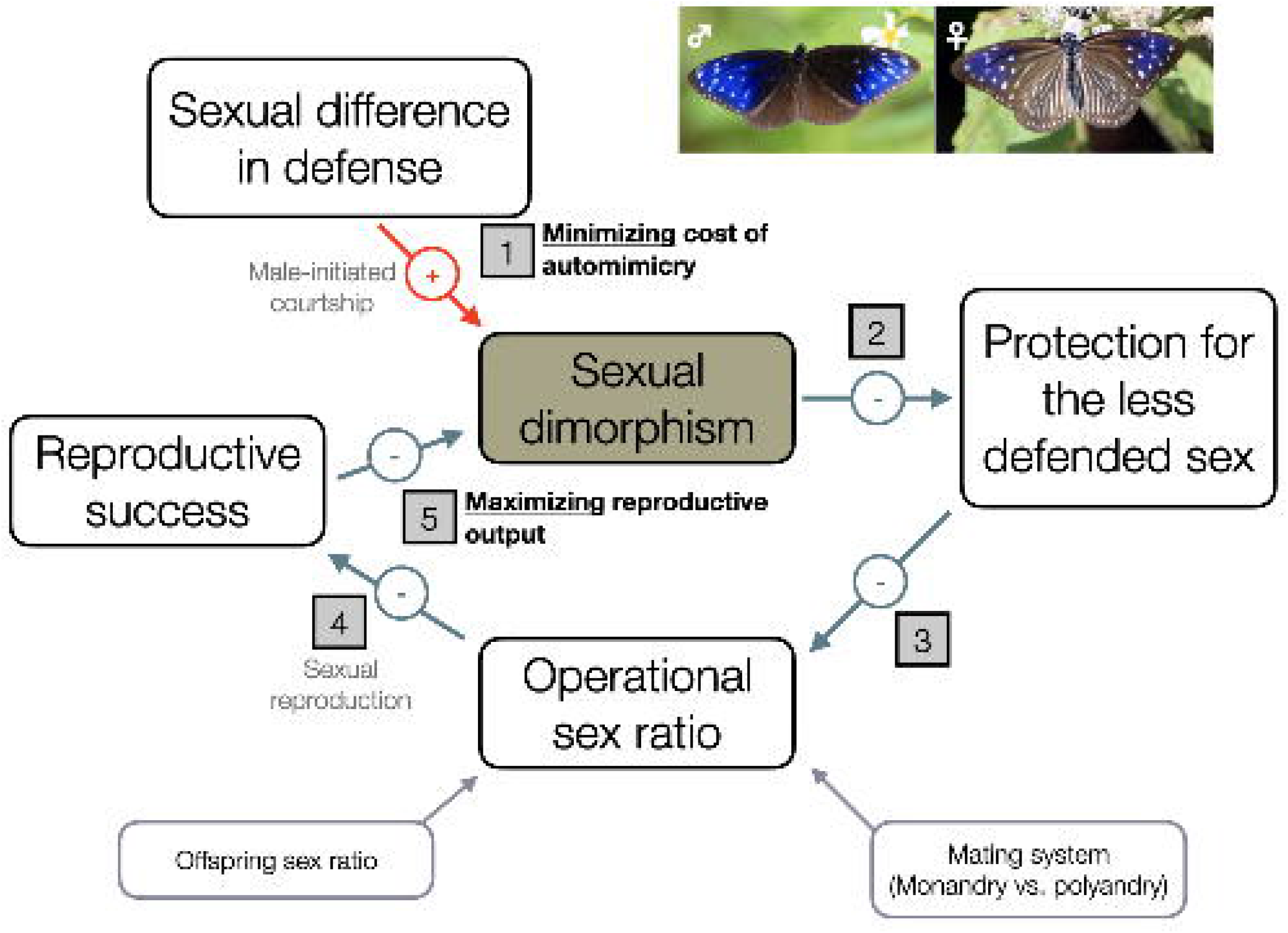
A conceptual diagram illistrating how minimizing the cost of automimicry and maximizing reproductive output can act in opposing directions and lead to the evolution of female-limited imprefect Müllerian mimicry in an aposematic species. The two photos show male (left) and female (right) *Euploea mulciber*. Photos were obtained from iNaturalist.

Here, I propose a hypothesis that intraspecific automimicry resulting from sexual difference in defense (automimicry, [19,20]) can lead to imperfect Müllerian mimicry in the less defended sex. The rationale behind this hypothesis rests on the balance between two antagonistic selective forces (Figure 1). On the one hand, the more defended sex may benefit from preferring members of the other sex with more dissimilar appearance, thereby reducing the cost of automimicry. The evolution of such mate preference could potentially drive the less defended sex to leave the Müllerian mimicry ring altogether (e.g., [17]). On the other hand, the loss of protection for the less defended sex as a consequence may be suboptimal in the context of reproduction, for which an even sex ratio would be ideal. It is therefore conceivable that the optimal appearance for the less defended sex could be a partial deviation from the mimetic pattern (i.e., imperfect Müllerian mimicry), with the extent of deviation determined by the degree of sexual difference in defense and the reproductive value of the less defended sex based on the operational sex ratio and mating system.

The abovementioned hypothesis could be particularly relevant in mimetic butterflies compared to other taxa for three reasons. First, males of numerous species employing chemical defense routinely replenish defense metabolites from diet in addition to their nutritious needs (pharmacophagy, [21]), resulting in high potential for sexual difference in defense. Second, many butterflies can often avoid hybridizing with comimics through chemical signaling (e.g., [22,23]). Selection for sexual dimorphism as a mechanism to reduce hybridization therefore might be weaker in those butterflies than species with less capability for chemical communication. Third, as courtship in butterflies is often initiated by males based on female color patterns [24], male mate preference has a higher potential for driving the evolution of female appearance. In addition, in some species females can only mate once in their lifetime (i.e., monandry), either due to the occurrence of mating plugs [25] or pupal mating [26]. In those monandrous species, the reproductive value of females will be much higher, thereby impacting the selective dynamics of female appearance and the degree of sexual dimorphism.

I use a modelling approach to test the hypothesis, with the model mechanics based on the general biology of butterflies (Figure 1). Specifically, I test two predictions. First, the antagonistic selection for minimizing the cost of automimicry and maximizing reproductive output can result in an optimal female appearance that is a partial deviation from the Müllerian mimetic pattern. Second, the degree of sexual dimorphism would be lower when females are reproductively more valuable either under a more male-biased offspring sex ratio or in a monandrous mating system.

## Material and Methods

### Model description

I modeled a Müllerian mimetic species in which males were more defended than females. This design was based on the biology of the focal species, but the principle was applicable to any species in which the sexes differed in defense and unprofitability. Since the focal species was part of a Müllerian mimicry ring, I assumed that signal appearance in males had fully converged with other comimics and did not vary. On the other hand, warning signal appearance could vary independently in females, such that deviation from mimicry in females, if favored by selection, did not affect mimicry fidelity in males. In this focal species, changes in the number of males and females can be described as the difference between births (*B*) and deaths (*D*):

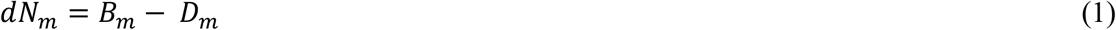

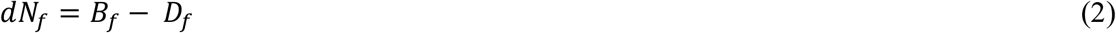

The increase in the number of males (*B*_*m*_) and females (*B*_*f*_) due to reproduction depended on whether females mated only once (monandry) or multiple times (polyandry) in their lifetime. In polyandrous species, increase in male and female numbers can then be described as:

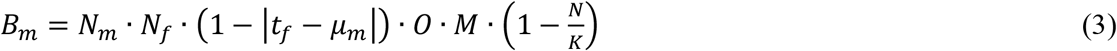

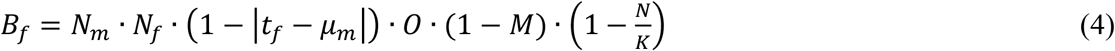

In monandrous species, however, the number of reproductive events was limited by the number of females, such that:

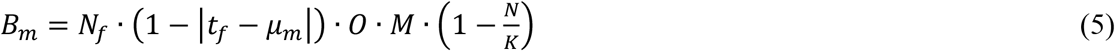

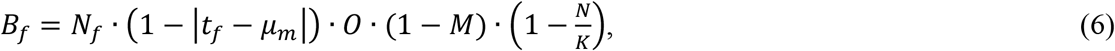

with *N*_*f*_ representing the number of females, *O* the number of offspring produced per reproductive event, *M* the proportion of sons produced (i.e., degree of male bias in the offspring sex ratio), and K the carrying capacity. Both *µ*_*m*_ and *t*_*f*_ take values between zero and one. The term *µ*_*m*_ signifies the female trait value most preferred by males. Female mate choice is not considered here, as male appearance does not vary. The term |*t*_*f*_ − *µ*_*m*_| thus describes the mismatch between female trait and male preference. The extent of mismatch, which also ranges from zero to one, determines the probability of mating. When |*t*_*f*_ − *µ*_*m*_| equals zero, female trait matches perfectly with male visual preference, resulting in maximal mating probability. Conversely, a |*t*_*f*_ − *µ*_*m*_| of one means female trait and male preference are completely mismatched, and no mating will occur. I set the difference between *t*_*f*_ and *µ*_*m*_ to be 0, 0.01, 0.03, 0.08, or 0.1 to examine the effect of matching male preference and female trait on the evolution of imperfect mimicry in females.

I considered predation to be the main cause of mortality for the system. Predation mortality for males (*D*_*m*_) and females (*D*_*f*_) can be described as the number of males and females multiplied by sex-specific predator attack rate:

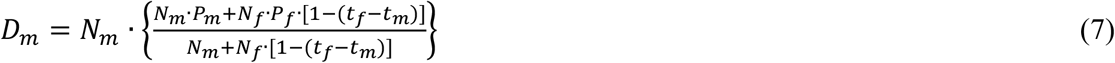

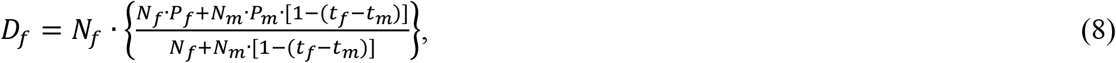

with *P* representing per capita predator attack rate, which differed between sexes if predators could distinguish between them (i.e., *P*_*m*_ ≠ *P*_*f*_), and *t* the visual appearance of the warning signal modelled in the context of predator’s foraging decision. In the context this study, predator attack rate was determined by the level of defense, with more defended sex having a lower value. I set *t*_*m*_ to 0 for mathematical convenience, whereas *t*_*f*_ can range from zero to one. When *t*_*m*_ equals *t*_*f*_, the two sexes were visually indistinguishable and the predator attack rates towards both sexes becomes the same weighted average 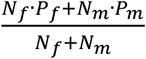. By contrast, when *t*_*f*_ equaled one, females were visually distinct from males, and there would be no cognitive generalization by predators. In this case, the per capita predator attack rate towards each sex reduced to *P*_*m*_ and *P*_*f*_, respectively.

### Model solution and fitness calculation

The species started from an initial population of 10 males and 10 females. To determine whether sexual difference in defense and biased offspring sex ratio might lead to sexual dimorphism in warning signal appearance, I chose 50 equidistant values between 0.01 to 0.4 for *P* and 50 values from 0 to 1 for *t*_*f*_. I excluded the parameter combinations in which *P*_*f*_ was lower than *P*_*m*_ as females should be less defended. I also chose ten equidistant *M* values between 0.1 and 0.9 to represent different levels of male bias in offspring sex ratio. I simultaneously solved equations (1) and (2) using the function *runsteady* in the R package *rootSolve* [27] for each parameter combination to find the number of males and females at equilibrium. Equilibrium population size (*N*_*m*_ + *N*_*f*_) for each *t*_*f*_ value under the same combination of *P*_*m*_, *P*_*f*_, and *M* was then scaled as the ratio to the maximum value to represent relative fitness.

## Results

The conditions in which imperfect mimicry was favored in females depended on mating system, offspring sex ratio, and the intensity of predation towards each sex (Figs 2 and 3). When females were monandrous and offspring sex ratio was unbiased, imperfect mimicry was only favored in females when sexual difference in defense exceeded a threshold (Fig 2A, B). As both sexes were attacked with higher frequencies, selection increasingly constrained the conditions under which imperfect mimicry was favored in females, as well as the degree of sexual dimorphism (Fig 2A-C). When offspring were already biased towards males, however, selection only favored imperfect mimicry in females when males experienced extremely low predation (Figs 2D-F and S1-S5); even when imperfect mimicry in females was favored, the extent of deviation from mimicry was more limited than that under an even offspring sex ratio (Figs 2A vs. 2D).

**Figure 2.**
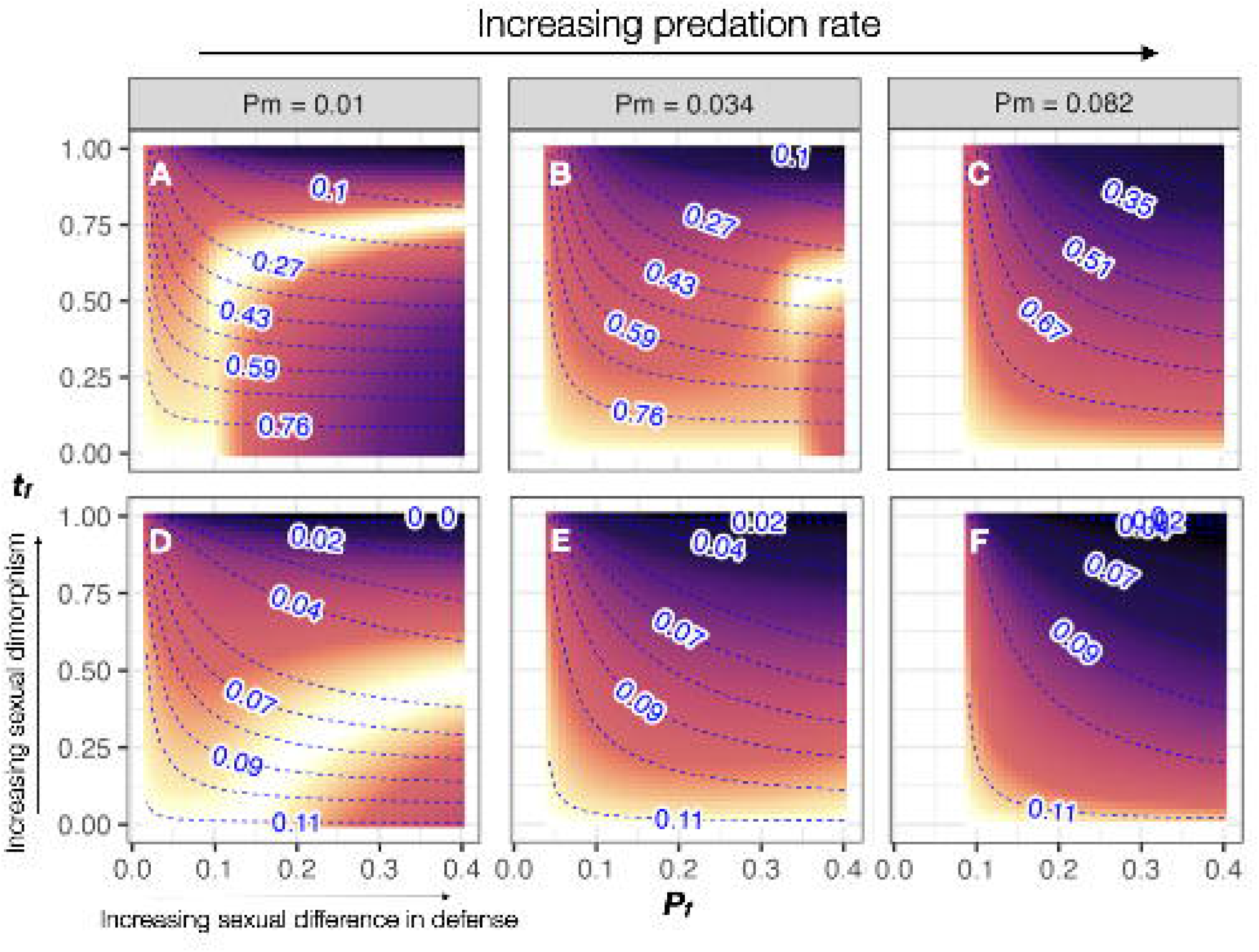
The degree of sexual dimorphism as favored by selection when offspring sex ratio is even (A-C) or extremely male biased (male:female ratio is 9:1, D-F) when females mate only once in their lifetime (monandry). Overall predation rate increases from left to right. In each subplot, larger values on the x-axis represent increasing sexual difference in defense, and larger values on the y-axis represent increasing sexual dimorphism. In each subplot, larger values on the x-axis represent increasing sexual difference in defense, and larger values on the y-axis represent increasing sexual dimorphism. Blue contours and numbers show the equilibrium female to male ratio. Brighter colors denote higher relative fitness.

**Figure 3.**
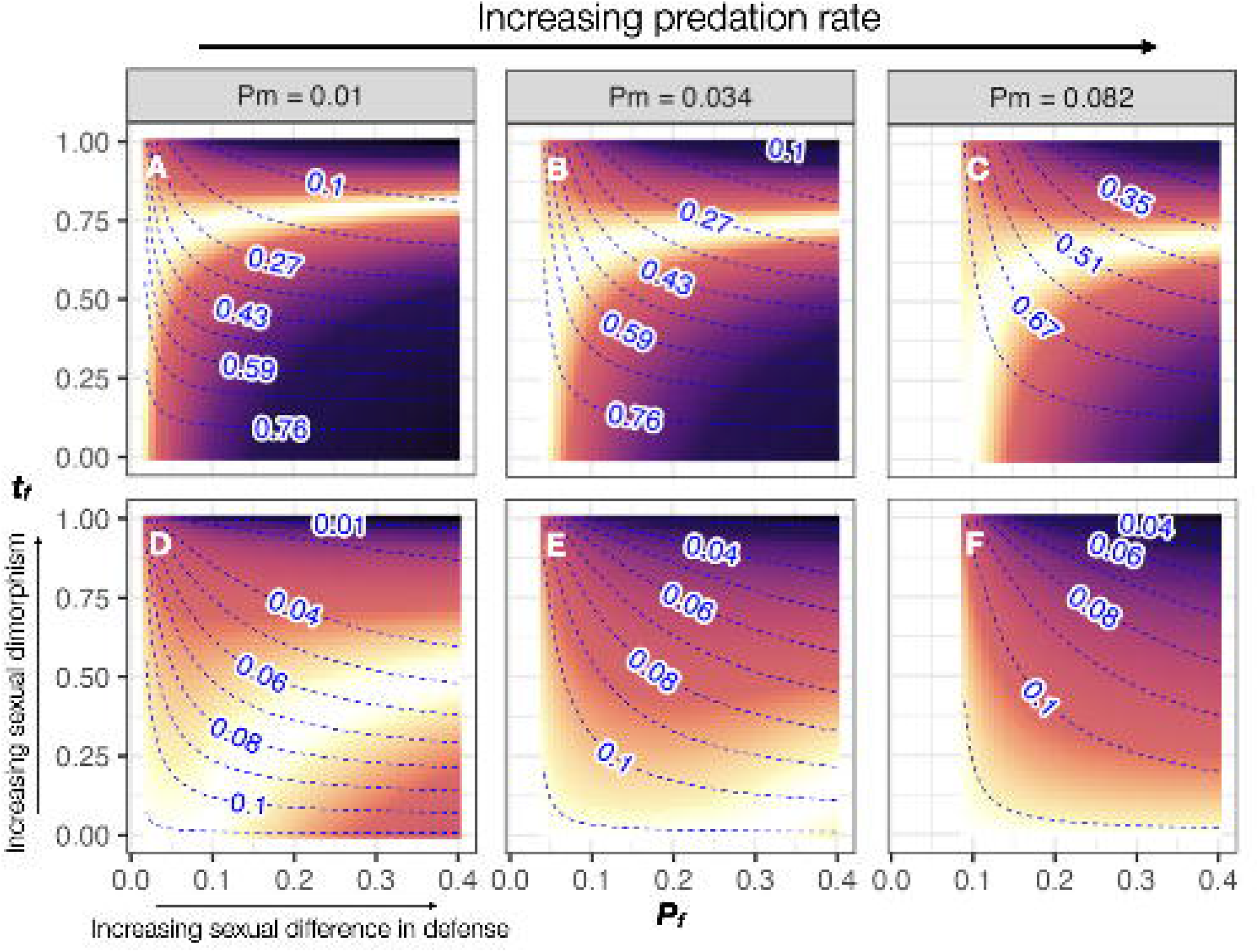
The degree of sexual dimorphism as favored by selection when offspring sex ratio is even (A-C) or extremely male biased (male:female ratio is 9:1, D-F) when females can mate multiple times throughout their lifetime (polynandry). Orientation of the plots and axis values on each subplot are the same as Figure 2. In each subplot, larger values on the x-axis represent increasing sexual difference in defense, and larger values on the y-axis represent increasing sexual dimorphism. Blue contours and numbers show the equilibrium female to male ratio. Brighter colors denote higher relative fitness.

When females were polyandrous and the offspring sex ratio was even, imperfect mimicry was favored as soon as there was slight sexual difference in defense, and increasing predation pressure did not severely prohibit the evolution of imperfect mimicry in females (Figs 3 and S6). Similar to the monandrous scenario, a male-biased offspring sex ratio further limited the evolution of sexual dimorphism, as well as the degree of sexual dimorphism when it was favored (Figs 3A-C vs. D-F, S6-S10). Notably, under no circumstance did selection favor females becoming completely nonmimetic, as a *t*_*f*_ equaling one invariably resulted in suboptimal relative fitness (Figures 2,3,S1-S10).

A mismatch between male visual preference and female trait only resulted in lower equilibrium population sizes but did not change the conditions under which imperfect mimicry was favored by selection in females (Table 1 and Figure S11). As courtship in butterflies are initiated by males, if selection favors male preference for females displaying an imperfect mimetic signal, female signal appearance is expected to evolve accordingly, and *vice versa*.

**Table 1.** Equilibrium population sizes under different *P*_*m*_ values and degrees of mismatch between male preference (*µ*_*m*_) and female trait (*t*_*f*_). Population sizes at equilibrium decreased when male preference and female trait were misaligned.

| $\vert t_f - \mu_m\vert $ | $P_m$ | | |
| --- | --- | --- | --- |
|  | 0.01 | 0.03 | 0.08 |
| 0 | 8919 | 8610 | 8179 |
| 0.01 | 8921 | 8603 | 8164 |
| 0.03 | 8911 | 8587 | 8136 |
| 0.08 | 8872 | 8542 | 8070 |
| 0.1 | 8846 | 8514 | 8032 |

## Discussion

Modeling results supported both predictions. Between minimizing the cost of automimicry and maximizing reproductive output, selection indeed favored partial departure from the common mimetic appearance in the less defended females, resulting in sex-limited imperfect Müllerian mimicry. Moreover, when females became more pivotal for reproduction – due to biased offspring sex ratio, a monandrous mating system, or simply higher overall predation rate – imperfect mimicry was favored in females under increasingly limited conditions, and the degree of possible sexual dimorphism was also reduced.

The modeling results offer insights into sexual dimorphism in aposematic species such as the butterfly *Euploea mulciber*, in which the sexes differ in defense and profitability (see Supplementary Materials). In *E. mulciber*, females have white lines and spots on the hindwings, which are absent in males and other *Euploea* comimics (Figure 1). This partial deviation from the mimetic pattern and the resulting female-limited imperfect mimicry could be driven by automimetic dynamics – a phenomenon that is relatively uncommon in aposematic species. Despite sexual difference in defense in *Euploea* butterflies, dimorphism in wing pattern only occurs in *E. mulciber*. Why female-limited imperfect mimicry has not evolved in other *Euploea* species requires further research, but the modeling results point to some possibilities. For example, it might be that offspring sex ratio is more male biased in the monomorphic species. Alternatively, even though there have not been reports of mating plugs or pupal mating in *Euploea* butterflies, females in the monomorphic species might be more limited in the number of matings they could participate during their lifetime. More detailed natural history data from these species would help test the plausibility of these possibilities.

The nature of sexual dimorphism in *E. mulciber* is distinct from that in *Oophaga* poison frogs and *Dilophotes* beetles. In *O. pumilio*, males are brighter in color than females, which is likely the product of Fisherian runaway selection due to directional female preference, but the two sexes do not differ in color pattern [12]. In *Dilophotes* beetles, the two sexes mimic different model species and bear little resemblance to each other (i.e., dual sex-limited mimicry) [10,11]. Even though it is currently unknown whether the two sexes of *Dilophotes* beetles differ in defense, findings from this study could potentially be relevant in explaining the evolution of such extreme sexual dimorphism if automimicry also occurs in these beetles. In a Müllerian system in which selection favors warning signal conformity, joining another mimicry ring inevitably requires deviating from current signal appearance and crossing an adaptive valley. However, the evolution of imperfect mimicry as an evolutionary solution to automimicry could represent a way to solve this conundrum and provide an intermediate step for extreme sexual dimorphism to evolve in Müllerian mimetic species. Even though dual sex-limited mimicry could lead to higher coextinction risk for the mimic and the model in Batesian systems [18], it should be ecologically more stable in Müllerian systems as all members are unprofitable.

In sum, this study presents a novel mechanism for the evolution of imperfect Müllerian mimicry in aposematic species in which one sex is consistently better defended than the other. Even though sexual dimorphism in Müllerian systems is rare, it nevertheless represents an important phenomenon that is counter to theoretical predictions. Through understanding this intriguing exception to the rule, we will be able to gain a more complete picture of how a multitude of selective forces might shape the diversity in prey phenotypes.

## Data accessibility

R codes and simulation data are available at: https://github.com/thekuolab/sex-limited-imperfect-Mullerian-mimicry

## Acknowledgements

The author thanks Hao-En Chin and Yu-Min Chung for their help with quantifying chemical defense in the *Euploea* butterflies.

## References

1. Müller F. 1879 Ituna and Thyridia; a remarkable case of mimicry in butterflies. (R. Meldola translation). Proclamations of the Entomological Society of London, 20–29.

2. Sherratt TN. 2008 The evolution of Müllerian mimicry. Die Naturwissenschaften 95, 681–695. (doi:10.1007/s00114-008-0403-y)

3. Mallet J, Joron M. 1999 Evolution of diversity in warning color and mimicry: polymorphisms, shifting balance, and speciation. Annual review of ecology and systematics 1999, 201–233. (doi:10.1146/ecolsys.1999.30.issue-1;wgroup:string:ar)

4. Walker H, Caro T, Bell D, Ferguson A, Stankowich T. 2023 Predation risk drives aposematic signal conformity. Evolution 77, 2492–2503. (doi:10.1093/evolut/qpad162)

5. Aubier TG, Sherratt TN. 2015 Diversity in Müllerian mimicry: The optimal predator sampling strategy explains both local and regional polymorphism in prey. Evolution 69, 2831–2845. (doi:10.1111/evo.12790)

6. Rowland HM, Wiley E, Ruxton GD, Mappes J, Speed MP. 2010 When more is less: the fitness consequences of predators attacking more unpalatable prey when more are presented. Biol Letters 6, 732–735. (doi:10.1098/rsbl.2010.0207)

7. Briolat ES, Gaston KJ, Bennie J, Rosenfeld EJ, Troscianko J. 2021 Artificial nighttime lighting impacts visual ecology links between flowers, pollinators and predators. Nat Commun 12, 4163. (doi:10.1038/s41467-021-24394-0)

8. Jamie GA, Dalziell AH, Welbergen JA, Tan EJ, Maguire C, Melamed E. 2025 The past and future of mimicry research. Nat. Ecol. Evol. 9, 1081–1085. (doi:10.1038/s41559-025-02775-8)

9. Kunte K, Kizhakke AG, Nawge V. 2021 Evolution of mimicry rings as a window into community dynamics. Annu Rev Ecol Evol Syst 52, 1–27. (doi:10.1146/annurev-ecolsys-012021-024616)

10. Motyka M, Bocek M, Kusy D, Bocak L. 2020 Interactions in multi-pattern Müllerian communities support origins of new patterns, false structures, imperfect resemblance and mimetic sexual dimorphism. Scientific Reports, 1–13. (doi:10.1038/s41598-020-68027-w)

11. Motyka M, Kampova L, Bocak L. 2018 Phylogeny and evolution of Müllerian mimicry in aposematic Dilophotes: evidence for advergence and size-constraints in evolution of mimetic sexual dimorphism. Sci. Rep. 8, 3744. (doi:10.1038/s41598-018-22155-6)

12. Maan ME, Cummings ME. 2009 Sexual dimorphism and directional sexual selection on aposematic signals in a poison frog. Proc. Natl. Acad. Sci. 106, 19072–19077. (doi:10.1073/pnas.0903327106)

13. Kunte K. 2008 Mimetic butterflies support Wallace’s model of sexual dimorphism. Proc. R. Soc. B: Biol. Sci. 275, 1617–1624. (doi:10.1098/rspb.2008.0171)

14. Su S, Lim M, Kunte K. 2015 Prey from the eyes of predators: color discriminability of aposematic and mimetic butterflies from an avian visual perspective. Evolution 69, 2985–2994. (doi:10.1111/evo.12800)

15. Long EC, Hahn TP, Shapiro AM. 2014 Variation in wing pattern and palatability in a female- limited polymorphic mimicry system. Ecol. Evol. 4, 4543–4552. (doi:10.1002/ece3.1308)

16. Wilson JS, Jahner JP, Forister ML, Sheehan ES, Williams KA, Pitts JP. 2015 North American velvet ants form one of the world’s largest known Müllerian mimicry complexes. Curr. Biol. 25, R704–R706. (doi:10.1016/j.cub.2015.06.053)

17. Maisonneuve L, Smadi C, Llaurens V. 2022 Evolutionary origins of sexual dimorphism: Lessons from female-limited mimicry in butterflies. Evolution 76, 2404–2423. (doi:10.1111/evo.14599)

18. Boutin M, Costa M, Fontaine C, Perrard A, Llaurens V. 2023 Influence of mimicry on extinction risk in Aculeata: a theoretical approach. Peer Community J. 3. (doi:10.24072/pcjournal.342)

19. Brower LP, Pough FH, Meck HR. 1970 Theoretical investigations of automimicry, i. Single trial learning. Proc. Natl. Acad. Sci. 66, 1059–1066. (doi:10.1073/pnas.66.4.1059)

20. Pough FH, Brower LP, Meck HR, Kessell SR. 1973 Theoretical Investigations of Automimicry: Multiple Trial Learning and the Palatability Spectrum*. Proc. Natl. Acad. Sci. 70, 2261–2265. (doi:10.1073/pnas.70.8.2261)

21. Boppré M, Monzón J. 2023 Baiting Insects with Pyrrolizidine Alkaloids (PAs): A Fieldwork-Oriented Review and Guide to PA-Pharmacophagy. Neotropical È ntomol. 52, 781–801. (doi:10.1007/s13744-023-01067-9)

22. González-Rojas MF, Darragh K, Robles J, Linares M, Schulz S, McMillan WO, Jiggins CD, Pardo-Diaz C, Salazar C. 2020 Chemical signals act as the main reproductive barrier between sister and mimetic Heliconius butterflies. Proceedings of the Royal Society B: Biological Sciences 287, 20200587. (doi:10.1098/rspb.2020.0587)

23. Darragh K et al. 2017 Male sex pheromone components in Heliconius butterflies released by the androconia affect female choice. PeerJ 5, e3953–23. (doi:10.7717/peerj.3953)

24. Cannon RJC. 2019 Courtship and Mating in Butterflies. Berkshire, SL5 7PY, UK: CAB International.

25. Carvalho APS, Orr AG, Kawahara AY, Carvalho APS, Kawahara AY. 2017 A review of the occurrence and diversity of the sphragis in butterflies (Lepidoptera, Papilionoidea). ZooKeys 694, 41–70. (doi:10.3897/zookeys.694.13097)

26. Walters JR, Corbin C, Hardcastle TJ, Jiggins CD. 2012 Evaluating female remating rates in light of spermatophore degradation in Heliconius butterflies: pupal-mating monandry versus adult-mating polyandry. Ecol. Èntomol. 37, 257–268. (doi:10.1111/j.1365-2311.2012.01360.x)

27. Soetaert K. 2009 rootSolve: Nonlinear root finding, equilibrium and steady-state analysis of ordinary differential equations. R package version 1.6. See http://CRAN.R-project.org/package=rootSolve.

