## Supplementary Materials for "Sexual difference in defense can drive the evolution of imperfect Müllerian mimicry in the less defended sex"

*Sexual difference in chemical defense in Euploea butterflies*

We collected five individuals of each sex from the following *Euploea* species during the summer of 2024 and 2025 in southern Taiwan (22°56'N 120°35'E): *E. tulliolus koxinga*, *E. sylvester swinhoei*, and *E. mulciber barsine*. We preserved all butterfly samples at -20 ºC immediately after collection. We pooled five individuals of the same sex and from the same year into one single sample for subsequent analysis. The defense chemical in this species is Intermedine-N-oxide, a pyrrolizidine alkaloid produced by numeroius plants to deter herbivores (Boppré, 1990; Boppré & Monzón, 2023). Prior to metabolomic analyses, we placed the samples in a freeze dryer for 24 hours before being weighed to the nearest 0.01 g, manually ground to fine powders, and then extracted with methanol. To prepare the analyte, we diluted the extract with methanol until a final concentration of 1000 ppm and then filtered through a 0.22 um PTFE filter. We quantified the amount of Intermedine N-oxide and its optic isomers using liquid chromatography coupled with triple quadruple mass spectrometry (LC-TQ-MSMS). We prepared a series of Intermedine N-oxide solutions with known concentrations (0, 5, 10, 25, 50, 80, and 150 ppb) from commercially available standard and used them to establish a calibration curve for interpolating the Intermedine N-oxide concentration in each sample. We calculated Intermedine N-oxide content per butterfly for each sex by multiplying the measured concentration by sample dry weight divided by five. All Euploea species showed clear sexual difference in the amount of Intermedine-N-oxide, with males being more chemically defended than females (Table S1).

Table S1. The amount of Intermedine N-oxide per individual in the two sexes of the three Euploea species. Numbers represent the average from samples in 2024 and 2025. Within each species, the first number is data from females, and the second number those from males.

|  | Species | | |
| --- | --- | --- | --- |
|  | E. tulliolus koxinga | E. mulciber barsine | E. sylvester swinhoei |
| Intermedine N-oxide (mg/individual) | 0.30, 0.57 | 0.08, 0.24 | 0.19, 0.80 |

**References**

Boppré, M. (1990). Lepidoptera and pyrrolizidine alkaloids Exemplification of complexity in chemical ecology. *Journal of Chemical Ecology*, *16*(1), 165–185. <https://doi.org/10.1007/bf01021277>

Boppré, M., & Monzón, J. (2023). Baiting Insects with Pyrrolizidine Alkaloids (PAs): A Fieldwork-Oriented Review and Guide to PA-Pharmacophagy. *Neotropical Entomology*, *52*(5), 781–801. <https://doi.org/10.1007/s13744-023-01067-9>

Soetaert, K. (2009). *rootSolve: Nonlinear root finding, equilibrium and steady-state analysis of ordinary differential equations. R package version 1.6*. <http://CRAN.R-project.org/package=rootSolve>

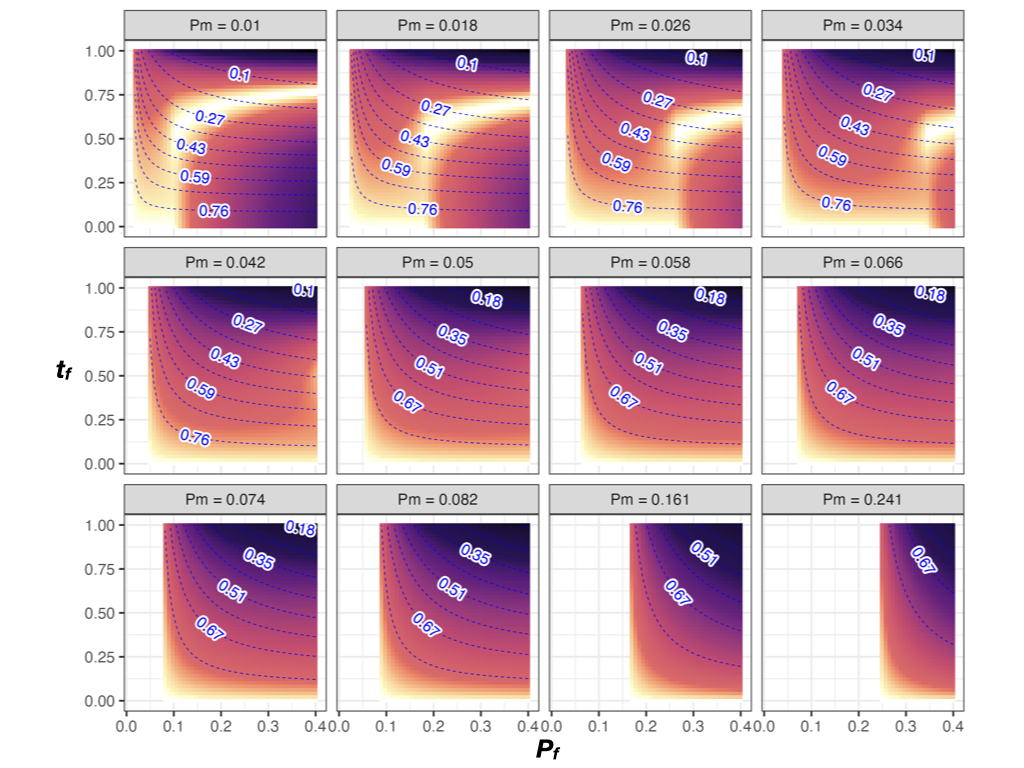

Figure S1. The degree of sexual dimorphism as favored by selection under a monandrous mating system and an even offspring sex ratio (M = 0.5). Overall predation rate increases from left to right and from top to bottom. In each subplot, larger values on the x-axis represent increasing sexual difference in defense, and larger values on the y-axis represent increasing sexual dimorphism. Brighter colors denote higher relative fitness. Blue contours and numbers are the equilibrium female to male ratio.

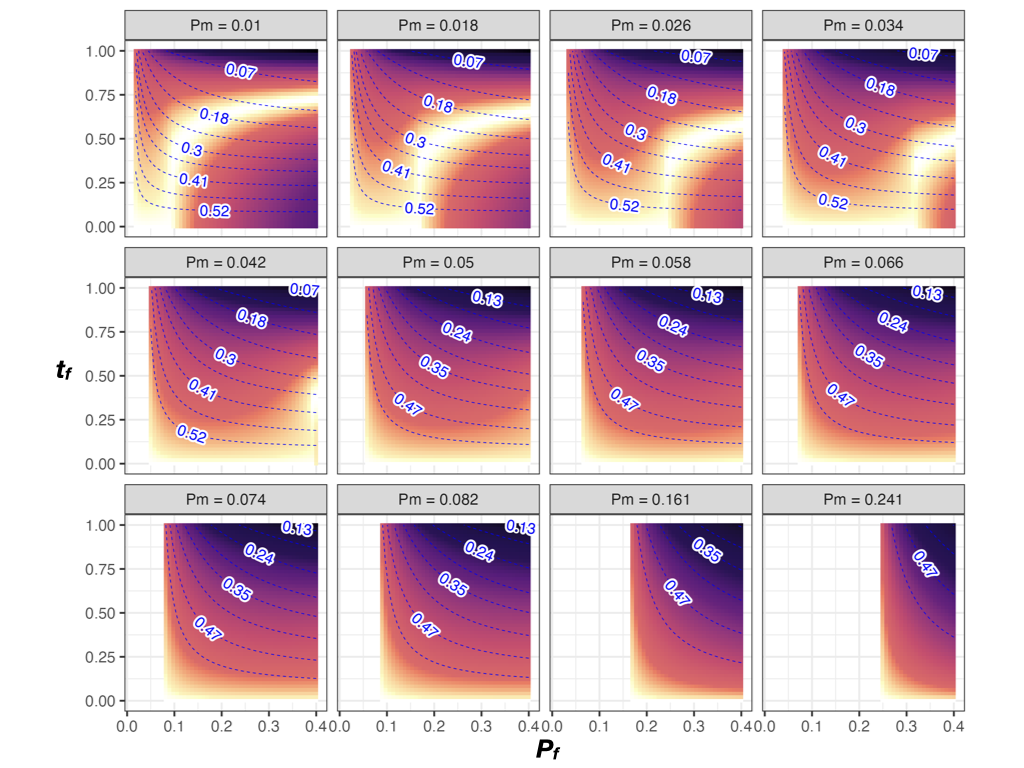

Figure S2. The degree of sexual dimorphism as favored by selection under a monandrous mating system and a slightly male-biased offspring sex ratio (M = 0.6). Overall predation rate increases from left to right and from top to bottom. In each subplot, larger values on the x-axis represent increasing sexual difference in defense, and larger values on the y-axis represent increasing sexual dimorphism. Brighter colors denote higher relative fitness. Blue contours and numbers are the equilibrium female to male ratio.

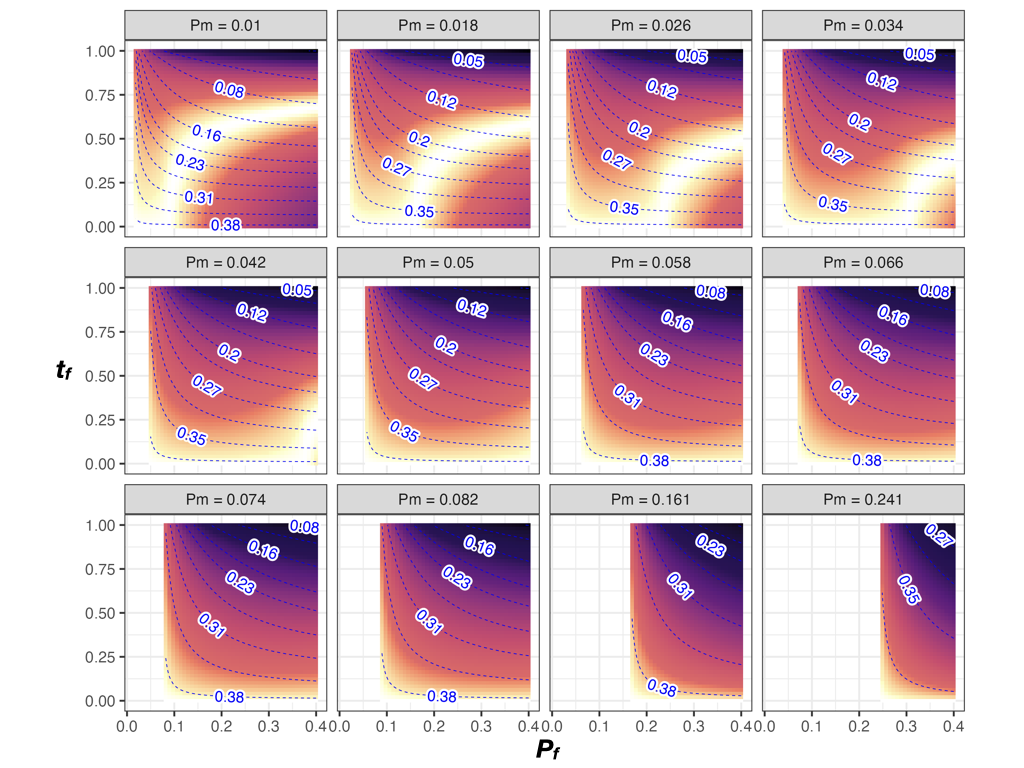

Figure S3. The degree of sexual dimorphism as favored by selection under a monandrous mating system and a moderately male-biased offspring sex ratio (M = 0.7). Overall predation rate increases from left to right and from top to bottom. In each subplot, larger values on the x-axis represent increasing sexual difference in defense, and larger values on the y-axis represent increasing sexual dimorphism. Brighter colors denote higher relative fitness. Blue contours and numbers are the equilibrium female to male ratio.

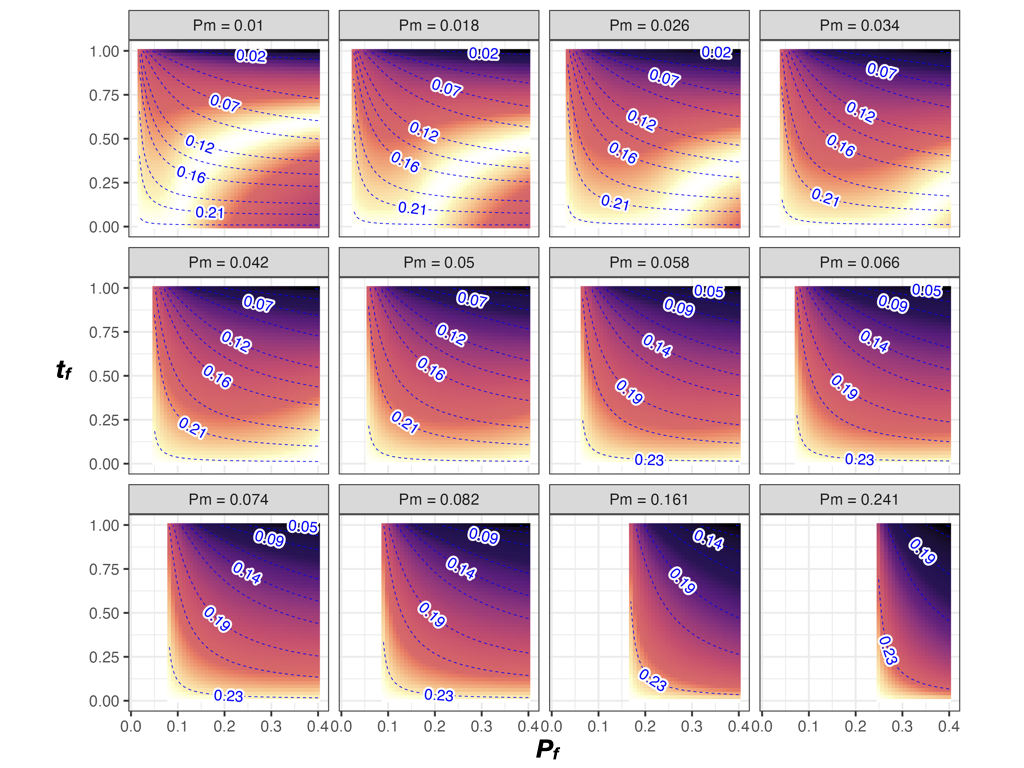

Figure S4. The degree of sexual dimorphism as favored by selection under a monandrous mating system and a highly male-biased offspring sex ratio (M = 0.8). Overall predation rate increases from left to right and from top to bottom. In each subplot, larger values on the x-axis represent increasing sexual difference in defense, and larger values on the y-axis represent increasing sexual dimorphism. Brighter colors denote higher relative fitness. Blue contours and numbers are the equilibrium female to male ratio.

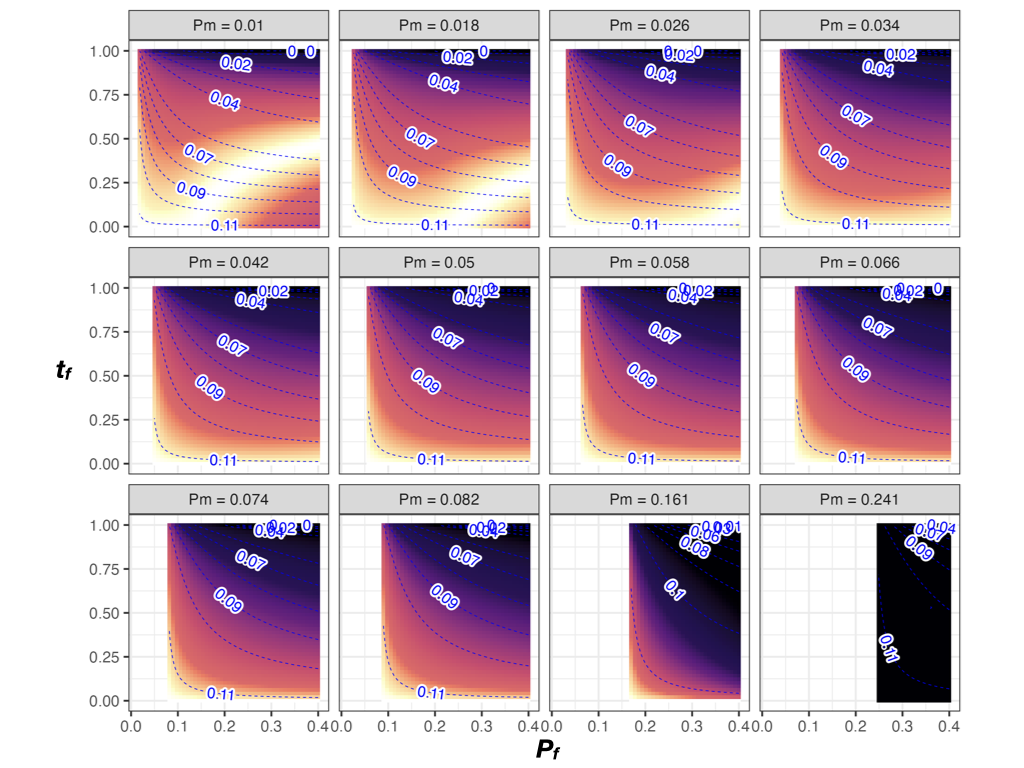

Figure S5. The degree of sexual dimorphism as favored by selection under a monandrous mating system and a extremely male-biased offspring sex ratio (M = 0.9). Overall predation rate increases from left to right and from top to bottom. In each subplot, larger values on the x-axis represent increasing sexual difference in defense, and larger values on the y-axis represent increasing sexual dimorphism. Brighter colors denote higher relative fitness. Blue contours and numbers are the equilibrium female to male ratio.

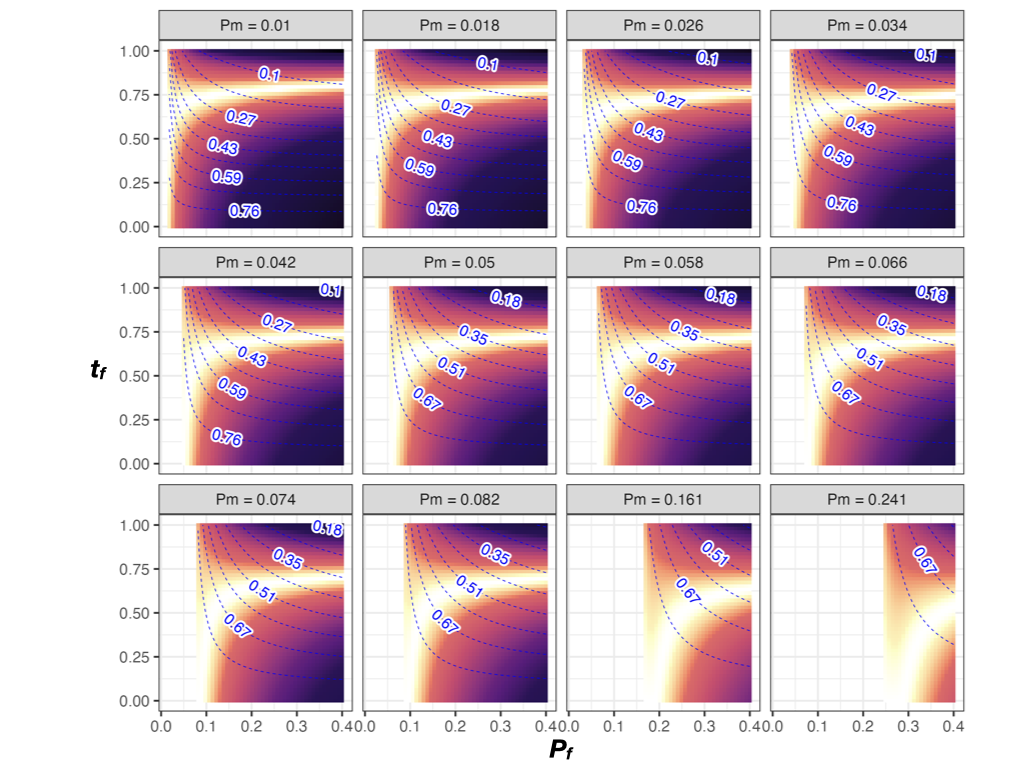

Figure S6. The degree of sexual dimorphism as favored by selection under a polyandrous mating system and an even male-biased offspring sex ratio (M = 0.5). Overall predation rate increases from left to right and from top to bottom. In each subplot, larger values on the x-axis represent increasing sexual difference in defense, and larger values on the y-axis represent increasing sexual dimorphism. Brighter colors denote higher relative fitness. Blue contours and numbers are the equilibrium female to male ratio.

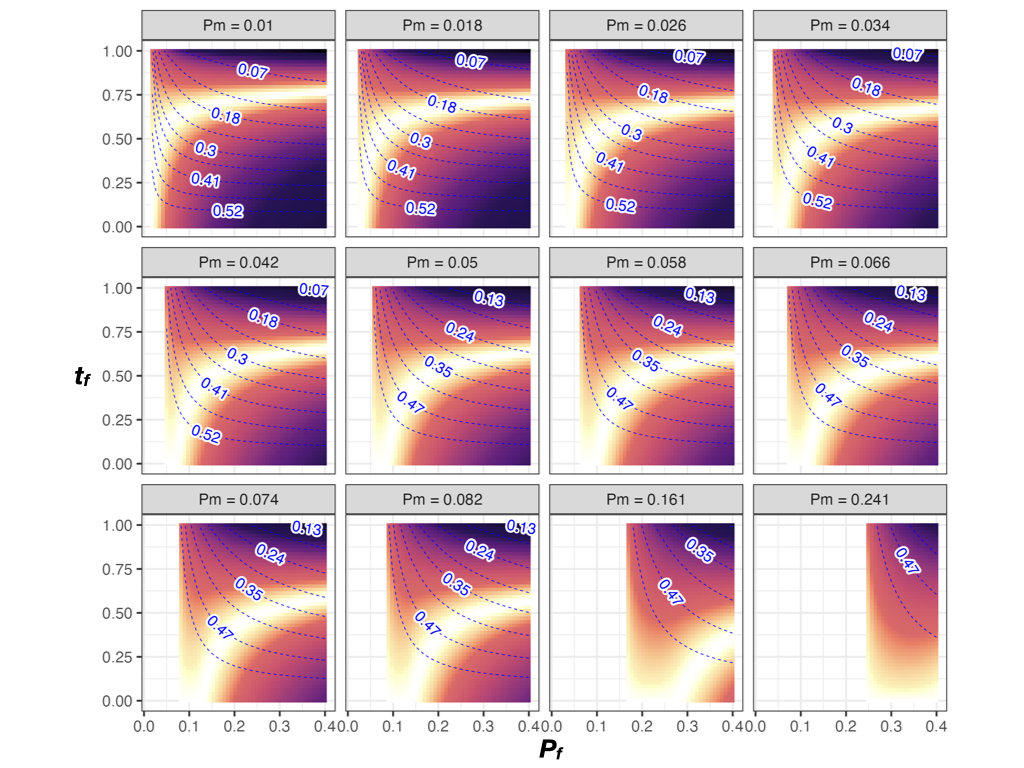

Figure S7. The degree of sexual dimorphism as favored by selection under a polyandrous mating system and a slightly male-biased offspring sex ratio (M = 0.6). Overall predation rate increases from left to right and from top to bottom. In each subplot, larger values on the x-axis represent increasing sexual difference in defense, and larger values on the y-axis represent increasing sexual dimorphism. Brighter colors denote higher relative fitness. Blue contours and numbers are the equilibrium female to male ratio.

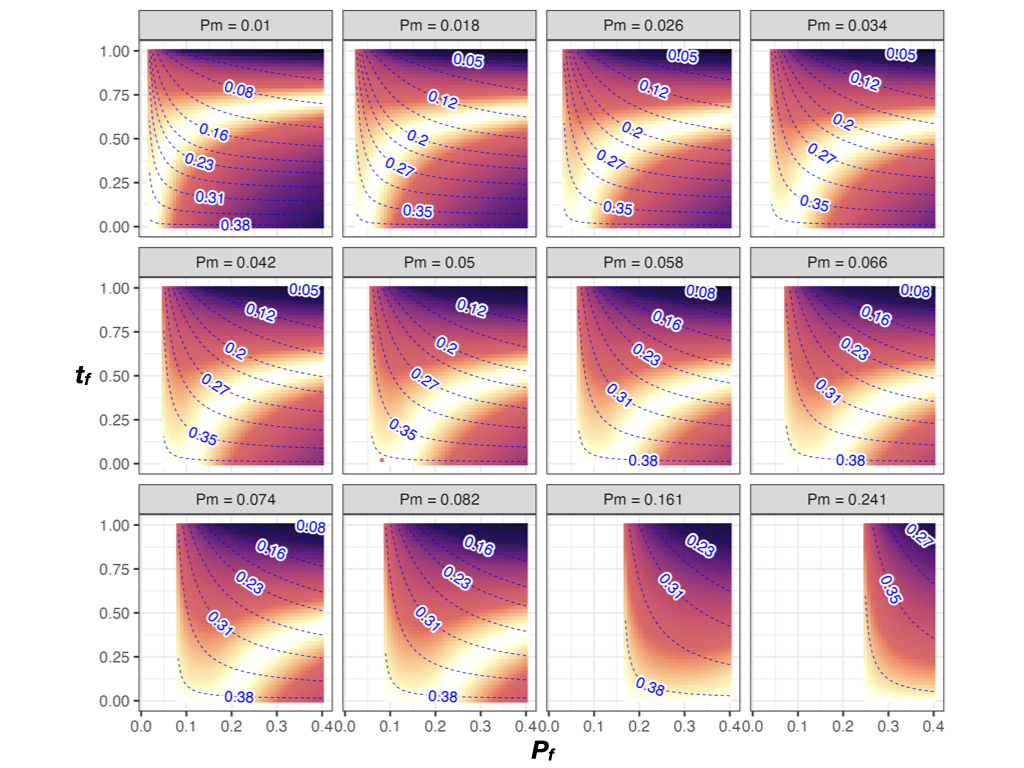

Figure S8. The degree of sexual dimorphism as favored by selection under a polyandrous mating system and a moderately male-biased offspring sex ratio (M = 0.7). Overall predation rate increases from left to right and from top to bottom. In each subplot, larger values on the x-axis represent increasing sexual difference in defense, and larger values on the y-axis represent increasing sexual dimorphism. Brighter colors denote higher relative fitness. Blue contours and numbers are the equilibrium female to male ratio.

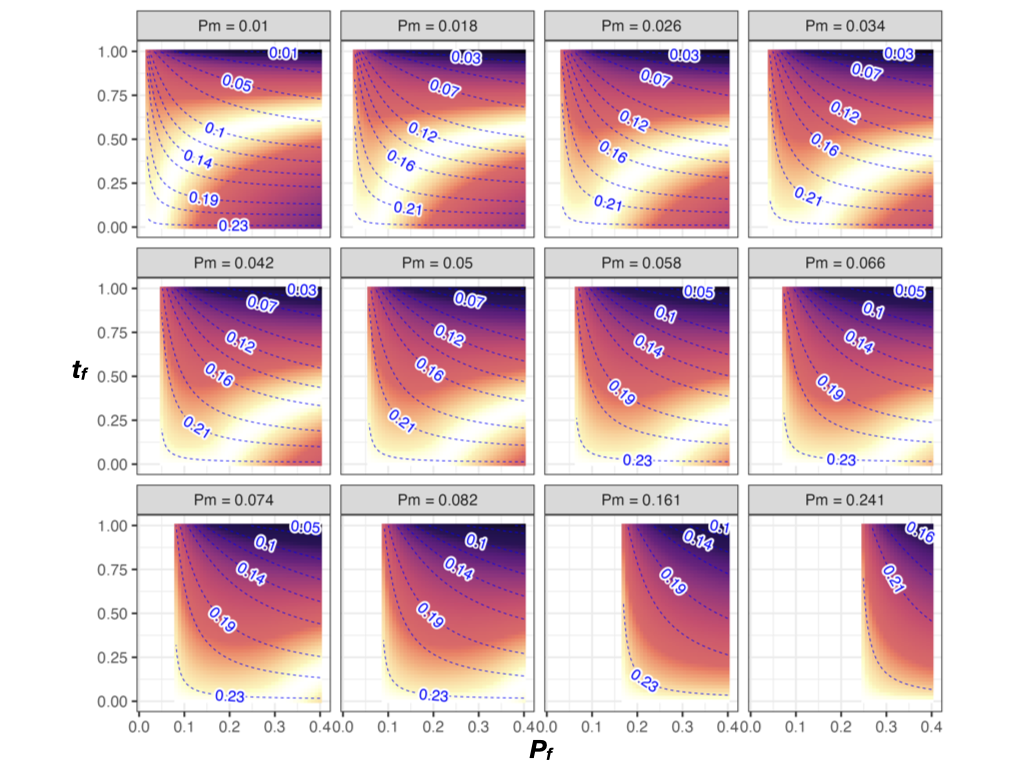

Figure S9. The degree of sexual dimorphism as favored by selection under a polyandrous mating system and a highly male-biased offspring sex ratio (M = 0.8). Overall predation rate increases from left to right and from top to bottom. In each subplot, larger values on the x-axis represent increasing sexual difference in defense, and larger values on the y-axis represent increasing sexual dimorphism. Brighter colors denote higher relative fitness. Blue contours and numbers are the equilibrium female to male ratio.

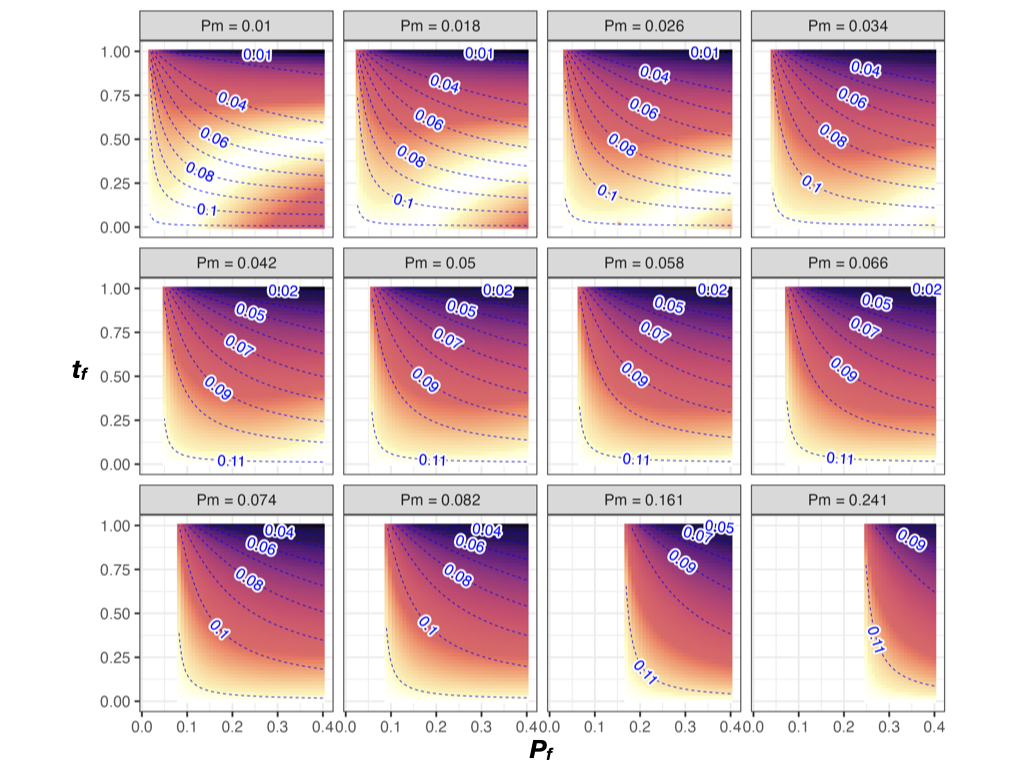

Figure S10. The degree of sexual dimorphism as favored by selection under a polyandrous mating system and a extremely male-biased offspring sex ratio (M = 0.9). Overall predation rate increases from left to right and from top to bottom. In each subplot, larger values on the x-axis represent increasing sexual difference in defense, and larger values on the y-axis represent increasing sexual dimorphism. Brighter colors denote higher relative fitness. Blue contours and numbers are the equilibrium female to male ratio.

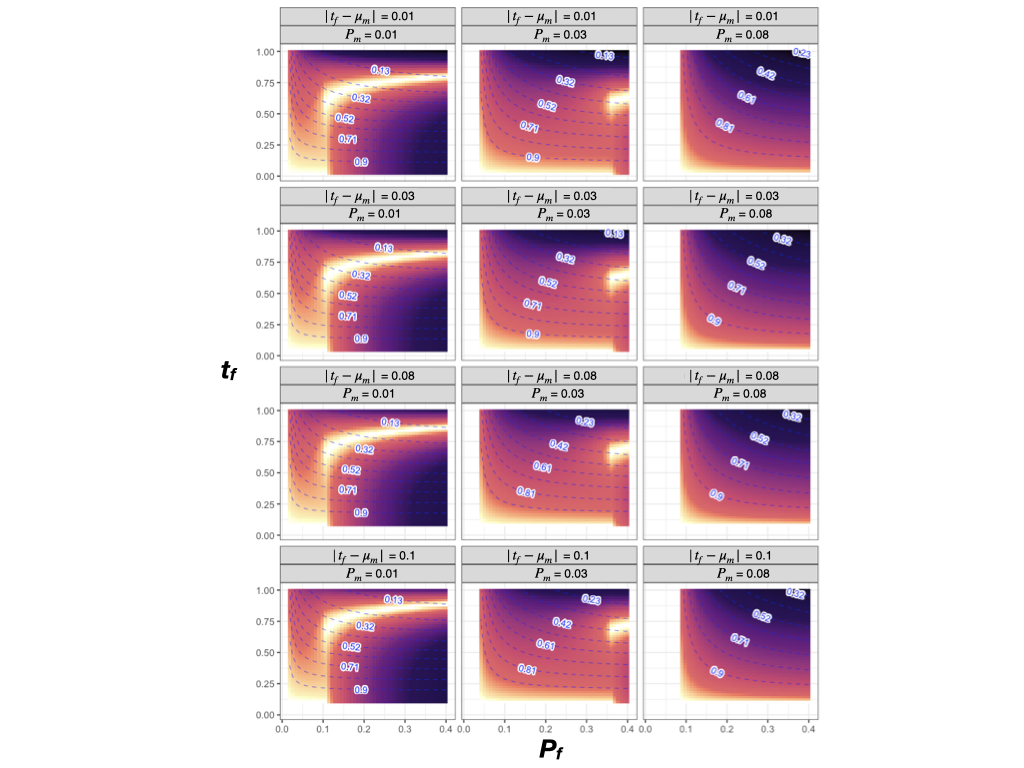
Figure S11. The degree of sexual dimorphism as favored by selection under increasing overall predation rate (*P_m_* = 0.01, 0.03, and 0.08) and increasing mismatches between male preference ($\mu_{m}$) and female trait ($t_{f}$) ($\left| t_{f}-\mu_{m} \right|$ = 0.01, 0.03, 0.08, and 0.1). In each subplot, larger values on the x-axis represent increasing sexual difference in defense, and larger values on the y-axis represent increasing sexual dimorphism. Brighter colors denote higher relative fitness. Blue contours and numbers are the equilibrium female to male ratio.
